# SARS-CoV-2 causes brain inflammation and induces Lewy body formation in macaques

**DOI:** 10.1101/2021.02.23.432474

**Authors:** Ingrid H.C.H.M. Philippens, Kinga P. Böszörményi, Jacqueline A. Wubben, Zahra C. Fagrouch, Nikki van Driel, Amber Q. Mayenburg, Diana Lozovagia, Eva Roos, Bernadette Schurink, Marianna Bugiani, Ronald E. Bontrop, Jinte Middeldorp, Willy M. Bogers, Lioe-Fee de Geus-Oei, Jan A.M. Langermans, Marieke A. Stammes, Babs E. Verstrepen, Ernst J. Verschoor

## Abstract

SARS-CoV-2 may cause acute respiratory disease, but the infection can also initiate neurological symptoms. Here we show that SARS-CoV-2 infection causes brain inflammation in the macaque model. An increased metabolic activity in the pituitary gland of two macaques was observed by longitudinal positron emission tomography-computed tomography (PET-CT). Post-mortem analysis demonstrated infiltration of T-cells and activated microglia in the brain, and viral RNA was detected in brain tissues from one animal. We observed Lewy bodies in brains of all rhesus macaques. These data emphasize the virus’ capability to induce neuropathology in this nonhuman primate model for SARS-CoV-2 infection. As in humans, Lewy body formation is an indication for the development of Parkinson’s disease, this data represents a warning for potential long-term neurological effects after SARS-CoV-2 infection.

**Teaser:** SARS-CoV-2 causes brain inflammation and Lewy bodies, a hallmark for Parkinson, after an asymptomatic infection in macaques.

## Introduction

Severe acute respiratory syndrome coronavirus 2 (SARS-CoV-2) causes a multi-system inflammatory disease syndrome, COVID-19 *(1)*. Although SARS-CoV-2 predominantly affects the respiratory organs, over 30% of the hospitalized COVID-19 patients also suffer from neurological manifestations, including loss of smell or taste, delirium, diminished consciousness, epilepsy, and psychosis *(2-5)*. Besides these general neurological symptoms, some patients additionally endure Parkinsonism *(6-8)*. The mechanisms behind this process are poorly understood. Neurological symptoms may be triggered by infection of the brain tissue, or indirectly, via virus-induced immune cell activation *(9)*. In humans, a direct link between brain inflammation and the presence of SARS-CoV-2 RNA has not been established yet *(10)*, and thus, formal proof that central nervous system (CNS)-related symptoms of COVID-19 are directly caused by the infection, or indirectly due to overactivation of the immune system, is lacking. Additionally, the long-term effects on the CNS after a mild to moderate SARS-CoV-2 infection, likely the vast majority of human cases, are unknown, and post-mortem brain samples from these individuals are not expected to become available for research in the near future. Controlled infection studies in a standardized experimental setting are crucial to investigate SARS-CoV-2-induced brain pathology *(11)*. To address this issue, a study was performed in two macaques species, rhesus macaques *(Macaca mulatta)* and cynomolgus macaques *(Macaca fascicularis)* (Table 1), both well-accepted animal models for COVID-19 *(12)*. Four male rhesus and four male cynomolgus macaques were inoculated with 10^5^ TCID_50_ of SARS-CoV-2 strain BetaCoV/BavPat1/2020 via a combined intratracheal and intranasal route *(13, 14)*. Following infection, SARS-CoV-2 genomic material was detected in tracheal and nasal swabs up to ten days, and based on clinical signs and thorax CTs, all animals showed mild to moderate disease symptoms *(13, 15)*.

**Table 1:**
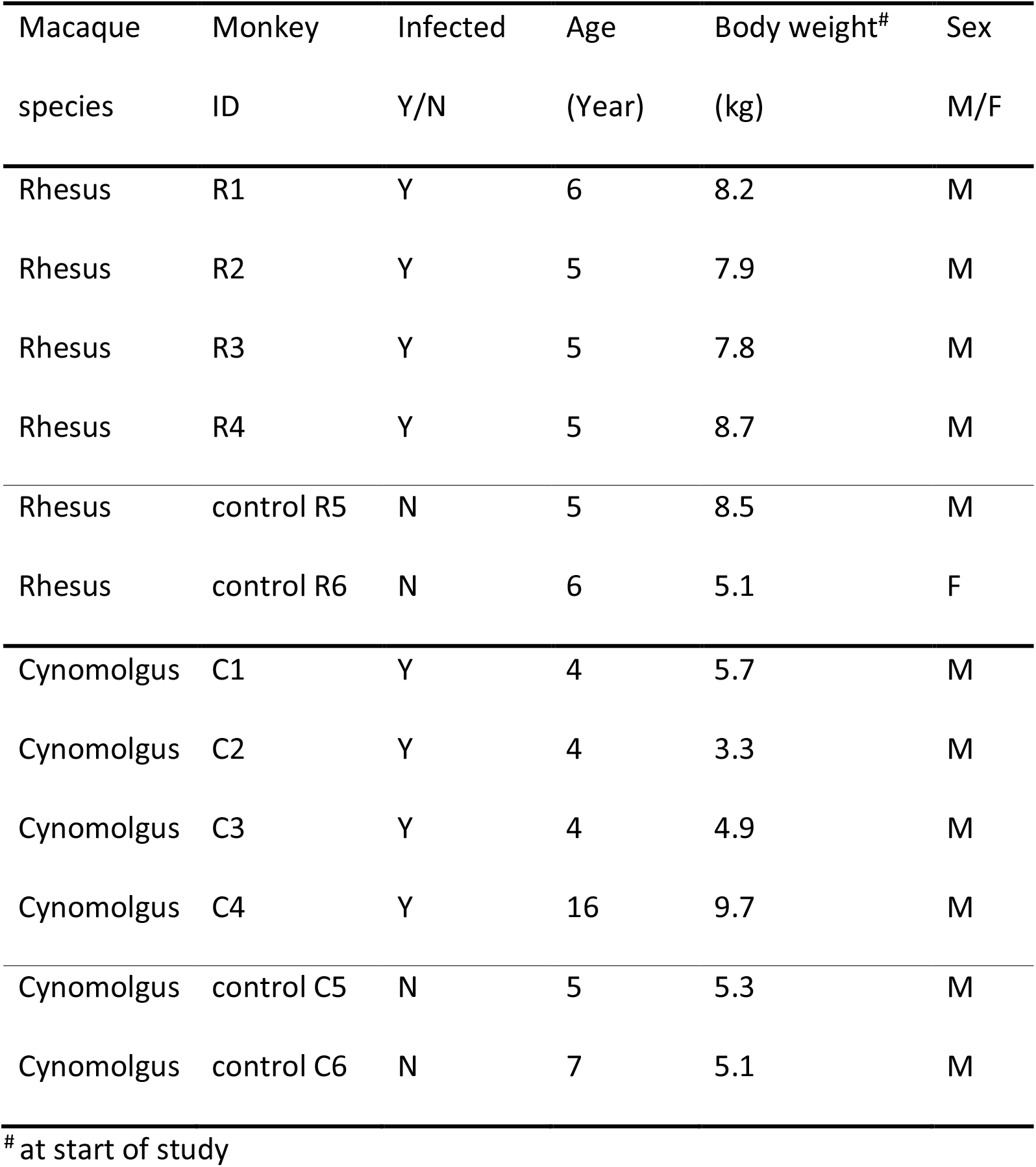
Animals featuring in this study.

## Results and Discussion

Weekly ^18^F-FDG PET-CTs of the brains of all macaques were initiated at the time point that the virus became undetectable in nasal and tracheal swabs. The uptake of tracer renders a marker for metabolic activity. Two of four cynomolgus macaques (C1 and C2) displayed an increased uptake of ^18^F-FDG in the pituitary gland at multiple time points. In animal C1, an increased uptake of ^18^F-FDG in the pituitary gland was seen at days 30 and 36 post-infection (Fig. 1). In animal C2, increased metabolic activity was already visible on day 8 and continued through day 35 when the final scan was obtained (Table S1).

**Fig. 1.**
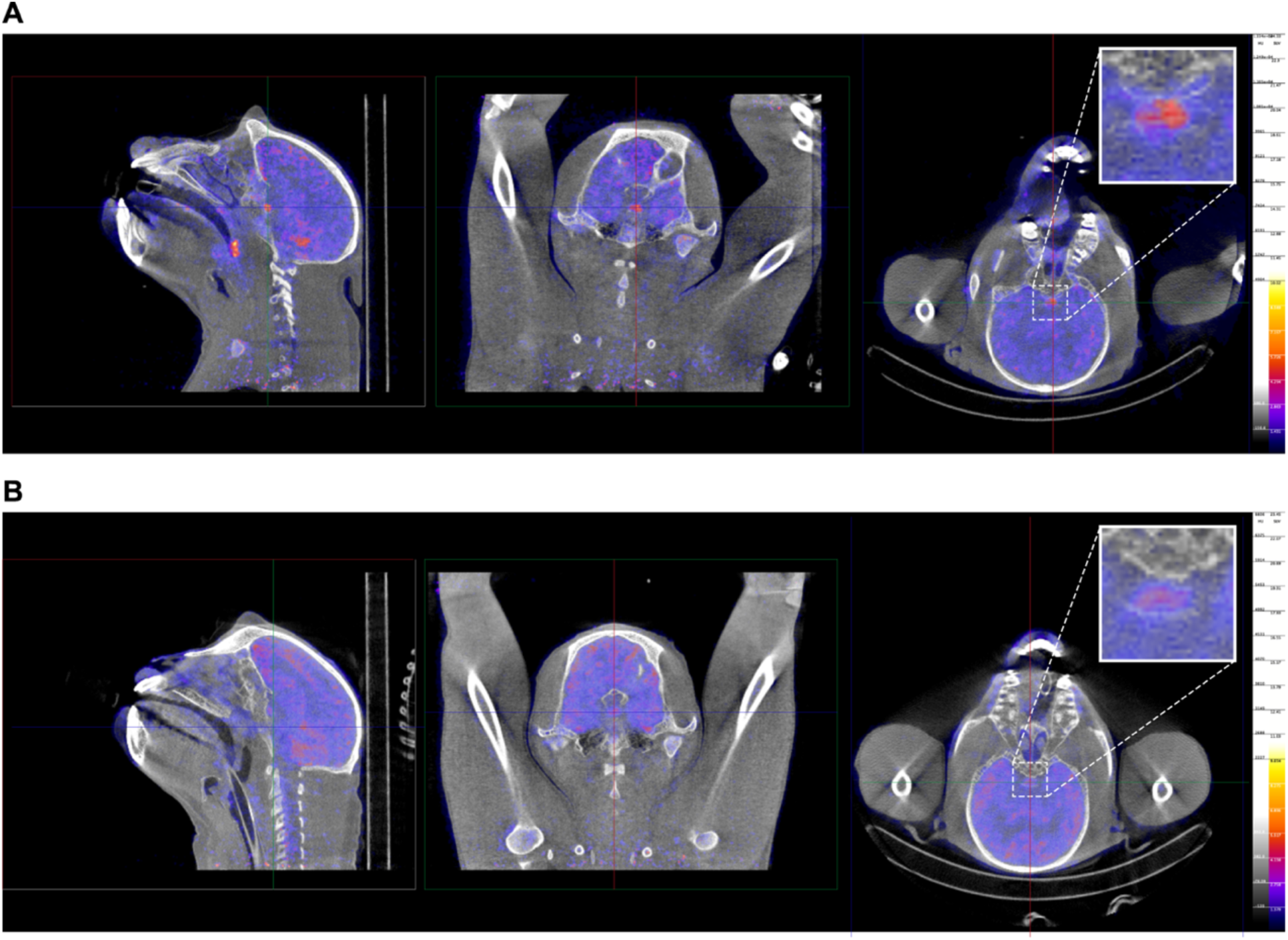
^18^F-FDG PET-CT image of a SARS-CoV-2 infected cynomolgus macaque (C1). Representative images of cynomolgus macaque C1 **(A)** on day 29, and a healthy control animal **(B)** are shown. The pituitary gland is indicated by the cross-hairs in all three directions and is boxed (right pictures). Similar window-level settings are applied for all sections. The pituitary gland-brain ratio of animal C1 was calculated with both the mean and peak standard uptake value (SUV); for the SUV_mean_ this ratio was 1.9, and for the SUV_peak_ 1.3. The average values calculated from a group of non-infected control animals (n=6) were 1.1 (std 0.3) for the SUV_mean_, and 0.6 (std 0.1) for the SUV_peak_ ratio. The uptake values of the brain, defined as background, were similar with SUV_mean_ values of 2.2 and 1.9 for the SARS-CoV-2-infected animals and the non-infected controls, respectively. After the relevant corrections for attenuation, scatter and decay, our results are indeed indicative for pituitary hypermetabolism after SARS-CoV-2 infection in animals C1 and C2.

In humans and macaques, the volume of the pituitary gland is small, and under physiological conditions, its metabolic activity is comparable to the background level of the entire brain *(16, 17)*. The ^18^F-FDG uptake of the pituitary gland may even be underestimated due to the partial-volume-effects that affect the emission signal recovery *(16)*. Because pituitary gland tissue expresses angiotensin-converting enzyme 2 (ACE2) *(18)*, the increased ^18^F-FDG uptake may be a direct effect of the infection or an indirect effect due to either a (reversible) hypophysitis, or transient hypothalamic-pituitary dysfunction *(19)*. Hypocortisolism has been reported as a delayed complication of SARS and has also been described in a SARS-CoV-2 patient *(20)*. For animal C1, which showed increased ^18^F-FDG uptake 30 days after infection, it is likely that the hypothalamic-pituitary axis was activated, leading to hypocortisolism similar to what has been found in patients with both a SARS and SARS-CoV-2 infection.

To further investigate the consequences of SARS-CoV-2-infection on macaque brain tissue, the animals were euthanized 5-6 weeks after experimental infection. Sections of the whole brain were systematically collected for further examination. As several regions of the brain express the SARS-CoV-2 receptor ACE2, and inflammation was found in the human brain *(10, 21, 22)*, we used various immunological markers for innate and adaptive immune activation to investigate for signs of immune activation, and also explored the localization of virus particles (Fig. 2 and Table 2). Viral RNA was detected by real-time quantitative polymerase chain reaction (RT-qPCR) in multiple brain areas of the right hemisphere of cynomolgus macaque C3 (Fig. 2). More precisely, cerebellum (1.48×10^5^RNA genome equivalents (GE)/gram), medial motor cortex (2.09×10^5^GE/gram), sensory cortex (2.07×10^5^GE/gram) and frontal basal cortex (8.29×10^4^GE/gram), as well as hippocampus (1.24×10^5^GE/gram), hypothalamus (1.05×10^6^ GE/gram), and globus pallidus (5.45×10^4^ GE/gram) all tested positive in the RT-qPCR. No viral RNA was detected in samples collected from the pituitary gland or olfactory bulb, substantia nigra, medulla oblongata, pons, nucleus caudatus, and putamen. Of interest, other tissues collected from macaque C3, including tracheobronchial lymph nodes, heart, liver, spleen, and kidney, also tested positive in the RT-qPCR, with comparable (lymph nodes, heart), or lower (kidney, liver, spleen) viral RNA loads *(13)*. Subgenomic messenger RNA analysis did not show evidence for active virus replication in the brains at the time point of euthanasia. Additionally, SARS-CoV-2 antigen was not detectable by immunohistochemistry the brains of all macaques.

**Table 2:**
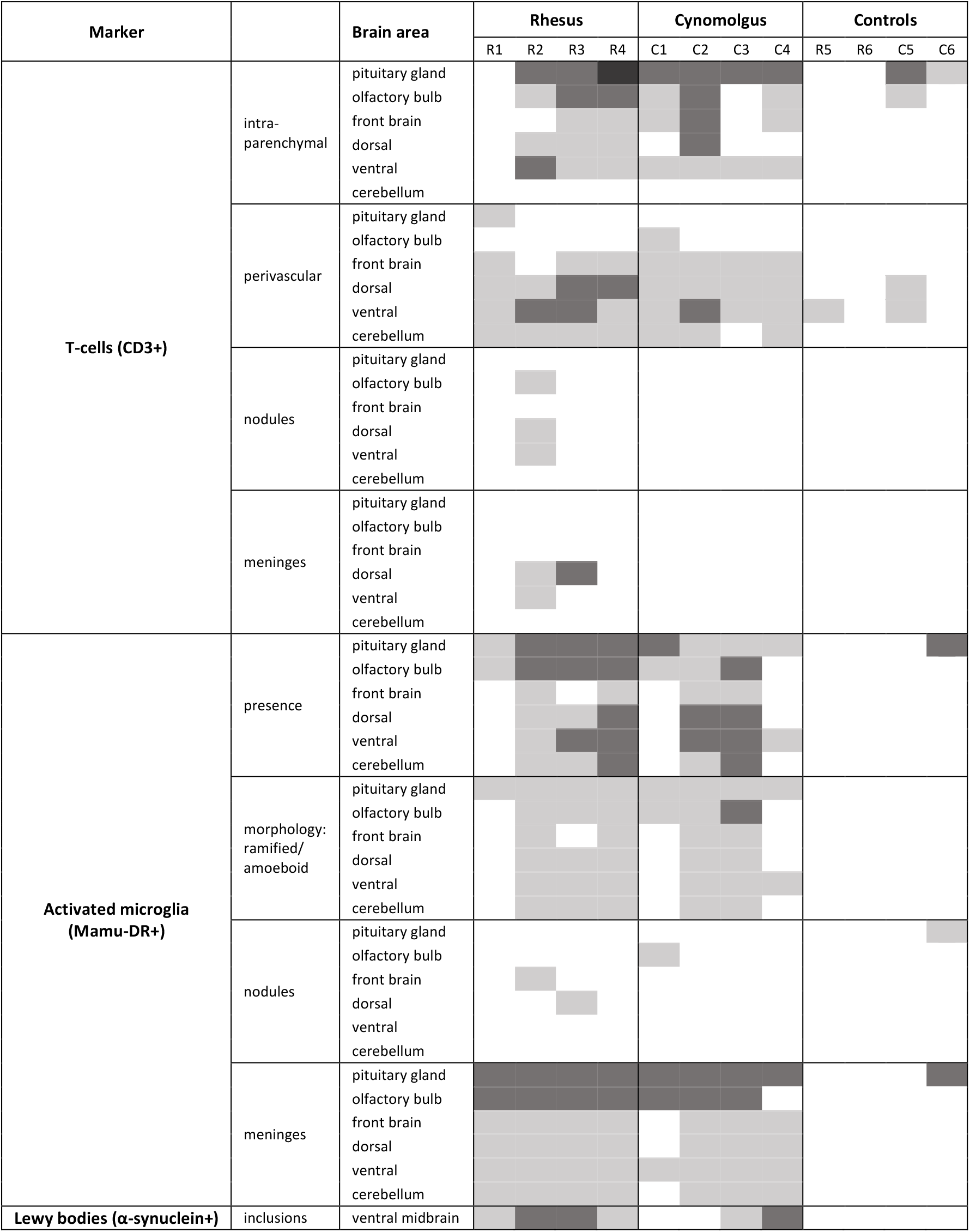
Histological findings. Table 2 outlines the presence of 1) T-cells (CD3+ cells) in the brain tissue (intraparenchymal), around blood vessels (perivascular), in group formation (nodules), or in the meninges, 2) activated microglia (Mamu-DR+ cells) in different parts of the brain, the morphology as a measure for the severity of activation (ramified or amoeboid), in group formation (nodules) or in the meninges, 3) α-synuclein/Lewy bodies (α-synuclein + cells with inclusions) in the ventral midbrain region next to the caudate nucleus. The last column shows the absence of most of these markers in the control animals. Light grey: mild observation; dark grey: moderate observation (including amoeboid microglia cells); black: moderate to severe observation of these markers.

**Fig. 2.**
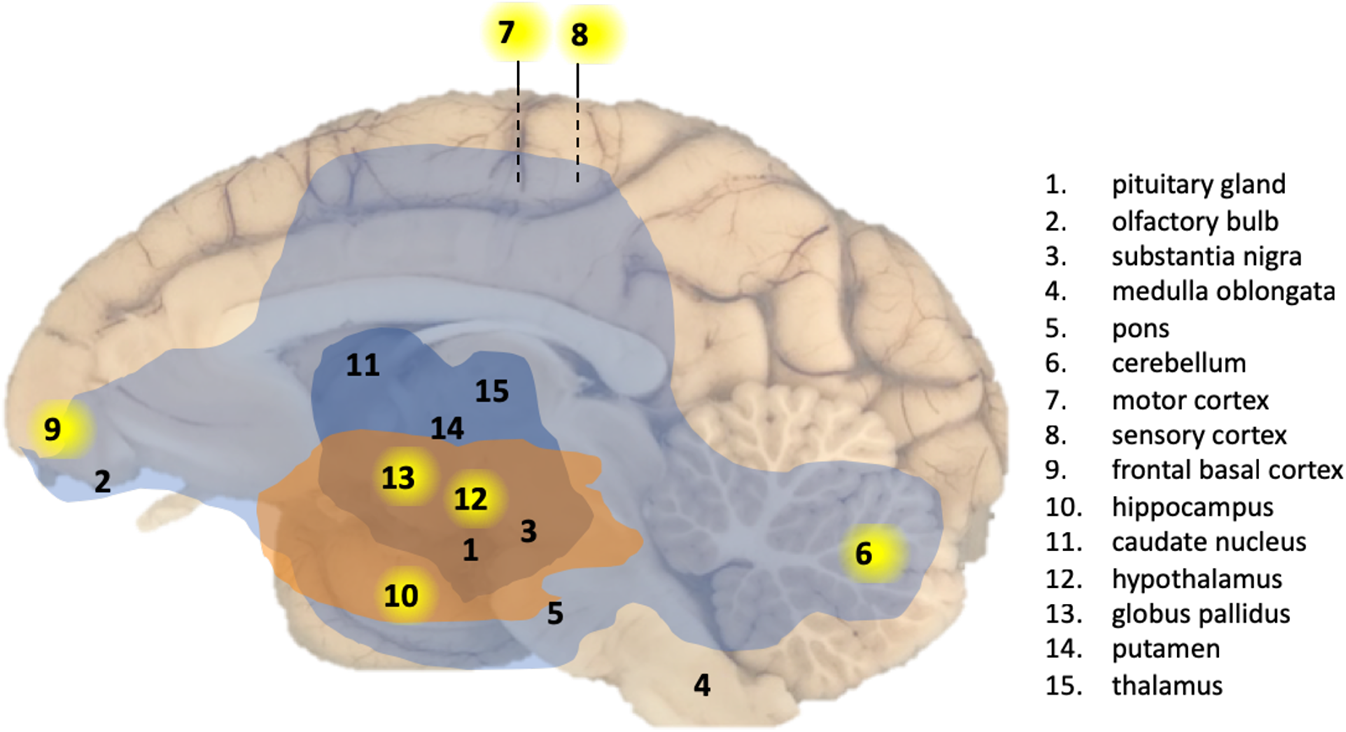
Overview of CNS effects by SARS-CoV-2 exposure in a macaque brain. Presence of viral RNA was investigated in multiple regions of the brain as indicated by the numbers. Viral RNA-positive regions in cynomolgus macaque C3 are indicated by a yellow background. The analysed brain regions are indicated with a number. Brain areas with T-cells (CD3+) and activated microglia (Mamu-DR+) are shown in light blue (mild expression) and dark blue (moderate expression), respectively. Brain areas with Lewy bodies (α-synuclein+) are shown in orange.

The brains of all SARS-CoV-2-infected macaques showed evidence of inflammation. Presence of T-cells was visualized by CD3 staining in intraparenchymal brain tissue, suggesting the infiltration of T-cells that passed the blood-brain barrier after SARS-CoV-2 infection (Fig. 3A, top panel). Additionally, activation of microglia cells in different areas of the brain, including the olfactory bulb and pituitary gland, was confirmed by Mamu-DR staining (upregulation of MHC class II expression) (Fig. 3A, middle panel). However, nodule formation, which is a measure for severity of activation, was rarely present (Table 2). No B-lymphocytic infiltration was found as evidenced by lack of CD20 staining (not shown). Hematoxylin and eosin (HE) staining did not show any abnormalities in the brain tissue of the virus-exposed macaques, including the absence of ischemic/necrotic lesions. For comparison, post-mortem brain tissues from two healthy, age-matched macaques of each species were used as controls (Table 1), none of the four control animals displayed obvious signs of immune activation (T-cells and microglia).

**Fig. 3.**
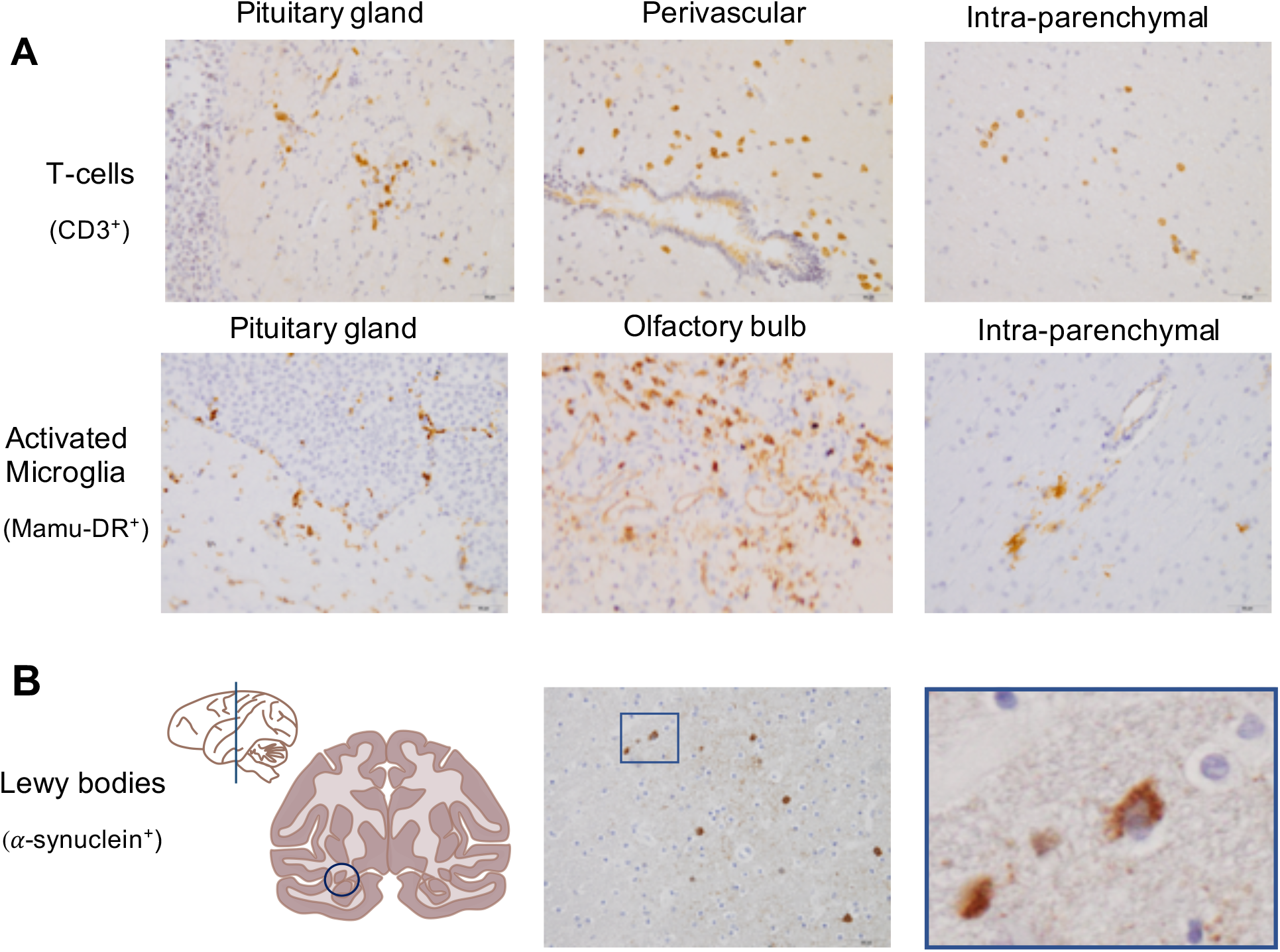
SARS-CoV-2 causes brain inflammation and Lewy body formationin brains of macaques. Immunohistochemical stainings of macaque brain tissues. **(A)** top panel: T-cells (CD3+) in the pituitary gland (left), perivascular (middle) and in the brain parenchyma (right) from a rhesus macaque (20x). **(A)** middle panel: activated microglia (Mamu-DR+) in the pituitary gland (left), olfactory bulb (middle) and in the brain parenchyma (right) from a cynomolgus macaque (20x). **(B)** bottom panel: Lewy bodies (α-synuclein +) were found in the ventral midbrain region next to the caudate nucleus indicated with a circle (left) in the coronal section of one hemisphere (anterior-posterior 0), presence of Lewy bodies in a SARS-CoV-2-infected rhesus macaque (20x) (middle), a magnified image (40x) of the square in the left image (right).

Brain tissues were screened for α-synuclein deposits, known as Lewy bodies, by immunohistochemistry. In humans, Lewy body formation is linked to the development of Parkinson’s disease or Lewy body dementia *(23, 24)*. It has been hypothesized that certain neurotropic viruses, including MERS and SARS coronaviruses, can trigger formation of Lewy bodies and cause Parkinsonism *(25-30)*.

The formation of intracellular Lewy bodies was clearly shown in the ventral midbrain region next to the caudate nucleus of all infected rhesus macaques (Table 2, Fig. 3B), and in one aged cynomolgus macaque (C4), while Lewy bodies were absent in the brains of all four control animals. For cynomolgus macaque C4, an age-dependent factor related to Parkinsonism cannot be excluded as this animal was older (16 years) than the other panel members (5-7 years), but the data from the rhesus macaques provide clear evidence for SARS-CoV-2-driven inflammation in the brain of macaques. In humans, neuropathology has been described in moribund COVID-19 patients, but we report of SARS-CoV-2-related brain involvement in macaques without displaying overt clinical signs. In general, macaque models for SARS-CoV-2 infection typically represent mild to moderate COVID-19 symptoms on the CT scan compared to humans *(12, 14, 31)*. Detection of viral RNA in the brain of an animal demonstrates the virus’ neuroinvasive capability. This matches a recent study describing neuroinvasion in mouse brains and in human brain organoids *(32)*.

How exactly SARS-CoV-2 caused widespread brain inflammation and induced Lewy body formation remains unknown. Viruses can enter the brain via different pathways. In this study, infiltration of T-cells was found perivascular and in the brain parenchyma, which indicates that the blood-brain barrier integrity may have been disturbed, offering the virus the opportunity to enter the brain. Alternatively, we hypothesize that SARS-CoV-2 gained access to the brain via neuronal pathways, such as the retrograde and anterograde neuronal transport through infected motor or sensory neurons *(33)*, and entered the pituitary gland via binding to the ACE2 receptor protein expressed on its cell surfaces *(18, 34)*. Such neural connection also exists between the olfactory bulb and the nasal mucosa *(35)*, and the loss of taste and smell, a characteristic of COVID-19, can thus be explained by nasal infection and subsequent inflammation in the olfactory bulb. Such a scenario is in line with the finding that in all the SARS-CoV-2-exposed macaques immune system activation in the olfactory bulb was evidenced by the presence of T-cells and/or activated microglia.

Neuronal transport can also explain why some COVID-19 patients develop Parkinson’s disease-like symptoms. Viruses can also, via retrograde transport in parasympathic motor neurons of the nervus vagus to the medulla, pons, and midbrain, reach the substantia nigra in the midbrain *(27, 36)*. Notably, the α-synuclein inclusions were found in the ventral midbrain region of the animals. In humans, these inclusions of accumulated misfolded proteins are associated with Parkinson’s disease or Lewy body dementia *(37)*.

There is a growing concern that symptomatic COVID-19 patients may suffer from long-term consequences *(9, 38)*. In this light the finding of Lewy bodies in brains of infected macaques without overt clinical signs is intriguing. Together with signs of inflammation and immune activation in the brains of the macaques this finding may point to a not yet earlier described SARS-CoV-2-induced neurodegenerative process that can explain the neurological symptoms that COVID-19 survivors experience *(39)*.

Lewy bodies are considered a hallmark for the development of Parkinson’s disease, or Lewy body dementia. More confirmation is required, but the observations in the translational macaque models for COVID-19 *(12-14, 40, 41)* can be regarded as a serious warning as they may be predictive for COVID-19-related dementia cases in humans in the future, even after an asymptomatic infection or mild disease process.

## Materials and Methods

### Animals and SARS-CoV-2 exposure

Four cynomolgus macaques *(Macaca fascicularis)* and four Indian-origin rhesus macaques *(Macaca mulatta)* (Table 1) were selected for this study. All macaques were mature, outbred animals, purpose-bred and socially housed at the BPRC. The animals were in good physical health with normal baseline biochemical and hematological values. All were pair-housed with a socially compatible cage-mate. The animals were offered a daily diet consisting of monkey food pellets (Ssniff, Soest, Germany) supplemented with vegetables and fruit. Enrichment was provided daily in the form of pieces of wood, mirrors, food puzzles, and a variety of other homemade or commercially available enrichment products. Drinking water was available *ad libitum* via an automatic system. Animal Care staff provided daily visual health checks before infection, and twice-daily after infection. The animals were monitored for appetite, general behavior, and stool consistency. All possible precautions were taken to ensure the welfare and to avoid any discomfort to the animals. All experimental interventions (intratracheal and intranasal infection, swabs, blood samplings, and PET-CTs) were performed under anesthesia. The research protocol was approved by national authorities (CCD, Central Committee for Animal Experiments; license number AVD5020020209404). Approval to start was obtained after further assessment of the detailed study protocol by the institutional animal welfare body (AWB) (in Dutch: Instantie voor Dierenwelzijn, IvD). The BPRC is accredited by the American Association for Accreditation of Laboratory Animal Care (AAALAC) International and is compliant with European directive 2010/63/EU as well as the “Standard for Humane Care and Use of Laboratory Animals by Foreign Institutions” provided by the Department of Health and Human Services of the US National Institutes of Health (NIH, identification number A5539-01).

On day 0, all animals were exposed to a dose of 10^5^ TCID_50_ of SARS-CoV-2 (strain BetaCOV/BavPat1/2020), diluted in 5 ml phosphate-buffered saline (PBS). The virus was inoculated via a combination of the intratracheal route (4.5 ml) and intranasal route (0.25 ml per nostril). For the histological examination brains from naive control macaques from the same age were obtained from the BPRC biobank, two cynomolgus, and two rhesus macaques.

### Positron Emission Tomography – Computed Tomography

Positron Emission Tomography (PET)-computed tomography (CT) data were acquired on multiple time points post-infection using a MultiScan Large Field of View Extreme Resolution Research Imager (LFER) 150 PET-CT (Mediso Medical Imaging Systems Ltd., Budapest, Hungary) as described before *(42)*. Animals were fasted overnight (glucose level > 8.5 mmol/l). The animals were sedated with ketamine (10 mg/kg ketamine hydrochloride (Alfasan Nederland BV, Woerden, The Netherlands)) combined with medetomidine hydrochloride (0.05 mg/kg (Sedastart; AST Farma B.V., Oudewater, The Netherlands)) to induce sedation and muscle relaxation, both applied intramuscularly (IM). The animals were positioned head first supine (HFS) with the arms up. The scans were acquired under mechanical ventilation in combination with a forced breathing pattern. For anesthetic maintenance, a minimum alveolar concentration of isoflurane (iso-MAC) of around 0,80%-1.00% was used. The body temperature of the animal was maintained by using the Bair Hugger (3M™, St Paul, MN, USA) supplied with 43°C airflow. Typically, around 100 MBq of ^18^F-FDG was applied intravenously (GE Healthcare, Leiderdorp, NL). Thirty minutes after injection the plateau in tracer activity uptake is reached, subsequently a PET of 15 minutes was acquired. After the scan, upon return to their home cage, atipamezole hydrochloride (Sedastop, ASTFarma B.V., Oudewater, NL, 5 mg/ml, 0.25 mg/kg) was administrated IM to antagonize medetomidine. Afterward the emission data was iteratively reconstructed (OSEM3D, 8 iterations, and 9 subsets) into a single frame PET image normalized and corrected for attenuation, scatter, and random coincidences using a reference CT and corrected for radioactive decay. The analysis was performed in VivoQuant 4.5 (Invicro, Boston, USA). Based on repeatability parameters for correct interpretation of the results, a standardized uptake value (SUV) ratio was used for robustness *(17, 42)*. An increased uptake, and pituitary gland hypermetabolism is defined as a SUV_mean_ ratio above 1.5 for the pituitary gland over the surrounding brain in combination with a SUV_peak_ ratio above background levels (>1.0). A group of non-infected control rhesus macaques (n=6) were used to calculate average background uptake of ^18^F-FDG.

### Brain tissue collection

Five weeks after virus exposure the macaques were euthanized and the brains were collected for further examination. The right hemisphere was used for RT-qPCR analysis and the left hemisphere was fixed in formalin for histology. Fifteen different regions were collected from the right hemisphere for RT-qPCR analysis: 1) part of the pituitary gland, 2) the olfactory bulb, 3) substantia nigra, 4) medulla oblongata, 5) pons, 6) anterior part of the cerebellum, 7) motor cortex medial, 8) sensory cortex, 9) frontal basal cortex, 10) hippocampus, 11) caudate nucleus, 12) hypothalamus, 13) globus pallidus, 14) putamen, and 15) thalamus. For the preparation of paraffin-embedded sections of the formalin-fixed left hemisphere, the cerebrum and cerebellum were dissected in 3-4 mm parts on the anterior-posterior axis. Pituitary gland and olfactory bulb were also embedded. From each part, sections (4 µm) were prepared for different staining methods. Immunohistochemistry stains were used for T-cells (CD3), B-cells (CD20), activated microglia (Mamu-DR), Lewy bodies (α-synuclein ab), and for SARS-CoV-2. Hematoxyline-eosine (HE) staining was used for general morphology.

### Viral RNA detection in brain tissue

Brain tissue samples were weighed and placed in gentleMACS M tubes (30 mg in 100 µl PBS) and treated using a gentleMACS Tissue Dissociator (protein01 program)(Miltenyi Biotec B.V., Leiden, The Netherlands). Next, the homogenized tissue was centrifuged, and 100 µl supernatant was used for RNA isolation. Viral RNA was isolated from using a QIAamp Viral RNA Mini kit (Qiagen Benelux BV, Venlo, The Netherlands) following the manufacturer’s instructions. Viral RNA was reverse-transcribed to cDNA using a Transcriptor First Strand cDNA Synthesis kit (Roche Diagnostics BV, Almere, The Netherlands). Viral RNA was quantified by RT-qPCR specific for RdRp gene of SARS-CoV-2, as described by Corman *et al*. *(43)*. The lower detection limit of the RT-qPCR was 3.6 viral RNA copies per reaction. Viral sub-genomic RNA was detected essentially as described by Wölfel *et al*. *(44)*. For both assays, RNA standard curves were generated by *in vitro* transcription of the target regions.

### Tissue preparation for histology

The left hemisphere of the brains, part of the pituitary gland, and one olfactory bulb were fixed in formalin for 24 hours and thereafter stored in buffered PBS. Preserved brains were cryoprotected in 30% w/v sucrose in PBS. The cerebrum was dissected in 12 different parts cut anterior-posterior axis at +10, +8, +5, +1, −3, −6, −8, −11, −14, −18, −22 from Bregma *(45)*, the cerebellum and pons were cut in 4 parts. These part were embedded in paraffin. From the eight brain parts in which viral RNA was detected by RT-qPCR, strips of brain sections were sliced into 12-series of 4 μm sections for different stains. These parts included the frontal cortex, midbrain parts, cerebellum, pituitary gland, and olfactory bulb. Sections were stained with virus antibody staining for virus detection and immunohistochemistry for immune reaction such as T-cell staining (CD3), B-cell staining (CD20), MHC-II cell staining (HLA-DR). Mirror sections were analyzed with a HE staining for brain morphology.

### Immunohistochemistry

The optimal concentration was determined for each antibody: CD3 (polyclonal rabbit – anti-human CD3 IgG, cat. no. A045201-2, Agilent Technologies), 1:60; CD20 (monoclonal mouse – anti-human CD20 IgG2a, clone L26, cat. no. M075501-2, Agilent Technologies), 1:800; Mamu-DR (monoclonal mouse – anti-human HLA-DR/DQ-IgG1, clone CR3/43, cat. no. M077501-2, Agilent Technologies), 1:150. For antigen retrieval, a steamer was used. Antigen Retrieval solution: IHC-TEK epitope retrieval solution, ready to use (catno IW-1100, IHC world). All incubation steps were at room temperature unless mentioned otherwise. Additionally, hematoxylin was used as a counterstaining in all protocols. After a dehydration sequence, the slides were mounted in Malinol. The counting of cells was performed in a blind matter.

#### CD3 and CD20 staining

The slides were deparaffinized by putting the slides sequentially in xylene, 100% ethanol, 96% ethanol, 70% ethanol, and PBS. Subsequently, an epitope antigen retrieval was executed in a steamer for 1h. After cooling down, the slides were placed in cuvettes (Sequenza cover plate system productnr 36107 Ted Pella inc.). Endogenous peroxidase (PO) activity was blocked by the PO blocking solution from DAKO (S2023) for 15 minutes. After a washing step (PBS with 0.05% Tween) Avidin was added from the DAKO kit (X0590) for 10 minutes. Thereafter, another washing step and biotin was added from the same DAKO kit (10 min) for blocking endogenous biotine. After washing a blocking step was executed for 20 minutes (PBS with 0.1% BSA and 1% normal human serum, NHS). The primary antibody was added (diluted in 0.1% BSA in PBS) and the slices were left overnight at 4°C. After washing a secondary antibody (Rabbit-anti-mouse IgG biotinylated (E0354), Agilent Technologies; 1:200 diluted in PBS + 1% BSA + 1% NHS) was added and, after washing, the slides were incubated with Vectastain ABC-peroxidase (ABC-PO, from Vector Laboratories; PK-4000; diluted 1:100 in PBS) for 30 minutes. After washing, 3,3’-diaminobenzidine (DAB) with 0.02% H_2_0_2_ was added to visualize the antigen-antibody binding (20 min).

#### Mamu-DR staining

The EnVision™ staining kit (G|2 Double-stain System, Rabbit/Mouse, DAB+/Permanent RED code K5361; Agilent technologies, Dako DK) was used for the immunohistochemical stain of Mamu-DR. The slides were deparaffinized by putting them sequentially in xylene, 100% ethanol, 96% ethanol, 70% ethanol, and PBS. Subsequently, an epitope antigen retrieval was executed and the slides were put in a steamer for 1 hour. The cooled down slides were placed in cuvettes and the endogenous peroxidase activity was blocked by the envision kit. The slides were washed and thereafter a blocking step was used consisting of 1% NHS + 1% BSA + 0.2% triton x100 in PBS for 10 minutes. Subsequently, the primary antibody was added (diluted in 0.1% BSA/PBS) for 30 minutes. Thereafter a washing step was implemented and the EnVision™ polymer/HRP (secondary antibody) was added for 10 minutes. Polymer HRP was added for 10 minutes followed by a washing step. Thereafter DAB+ was added for 15 minutes to visualize the antigen-antibody binding.

#### α-Synuclein staining

The slides were deparaffinized by immersing them sequentially in xylene, 100% ethanol, 96% ethanol, 70% ethanol, and PBS. Subsequently, an epitope antigen retrieval was executed by putting the slides for 15 minutes in Formic acid (100%) diluted 1:10 in demineralized water. After 2 washing steps in PBS with 0,05% Tween, the slides were placed in cuvettes (Sequenza cover plate system product no. 36107 Ted Pella inc.). Endogenous PO activity was blocked by the PO blocking solution from DAKO (S2023) for 20 minutes. After washing (PBS with 0.05% Tween), avidin was added from the DAKO kit (X0590) for 10 minutes. Then, after another washing step, biotin was added from the same DAKO kit (10 min) for blocking of endogenous biotine. After washing, a blocking step was executed for 30 minutes (PBS with 0.1% BSA and 1% NHS and 0.02% Triton-X100). The primary antibody, α-synuclein clone 4D6 (Biolegend SIG-39720), was added (diluted in 0.1% BSA in PBS) and the slides were left overnight at 4°C. After washing, a secondary antibody (rabbit anti-mouse IgG Biotinylated (E0354), Agilent Technologies; 1:200 diluted in PBS+ 1% BSA) was added for 30 minutes and the slides were incubated with Vectastain ABC-PO kit from Vector Laboraties (PK-4000; diluted 1:100 in PBS) for 30 minutes. After a final washing step, DAB with 0.02% H_2_0_2_ was added to visualize the Antigen-antibody binding (20 min).

#### SARS-CoV-2 staining

The Roche Optiview DAB IHC kit was used in a Ventana Benchmark Ultra (Roche, Basel Switzerland) immunostainer to immunohistochemically stain SARS-CoV-2. Two monoclonal antibodies of ThermoFisher raised to SARS-CoV-2 Nucleocapsid (clone B46F, catno MA1-7404, and E16C, catno. MA1-7403) were validated on formaldehyde-fixed and paraffin-embedded SARS-CoV-2 and mock-infected Vero E6 cells *(10)*, as well as lung tissue sections of human SARS-CoV-2 patients. The clone E16C was superior to B46F and was further used in this study. Antigen retrieval took place with cell conditioning 1 (CC1, Ventana Medical Systems) (pH 8,5) for 24 minutes at 100°C, 1/5.000 diluted. Thereafter, incubation took place with the primary antibody for 48 minutes at 36°C followed by standard Optiview detection/visualization with DAB and Copper. After immunohistochemical staining, the sections were dehydrated with grades of ethanol and cleared with xylene. All sections were mounted with TissueTek® coverslipping film (Sakura Finetek Europe B.V., Alphen aan den Rijn, The Netherlands).

## Acknowledgements

We want to thank Francisca van Hassel for assistance with editing of figures, Yolanda Kap for reviewing the manuscript, Wim Vos for the excellent histology, and the Animal Science Department of the BPRC, the veterinarians and animal caretakers for all the experimental support.

## Author contributions

Conceptualization: IHCHMP, EJV, MAS, BEV

Data curation: MAS, BEV, IHCHMP

Formal Analysis: MAS, IHCHMP, BEV

Funding acquisition: EJV

Investigation: KPB, JAW, ZCF, NvD, AQM, DL

Methodology: MAS, MB, BS, IHCHMP

Project administration: EJV, BEV

Resources: REB

Supervision: IHCHMP, MAS, BEV, EJV

Validation: MB, ER, L-FG-O,

Writing – original draft: IHCHMP, KPB

Writing – review & editing: MAS, EJV, BEV, JAML, REB, JM, WMB, L-FG-O

## Data and materials availability

All data needed to evaluate the conclusions in the paper are present in the paper and/or the Supplementary Materials. Correspondence should be addressed to IHCHMP.

## Supplementary material

**Supplementary Table S1:**
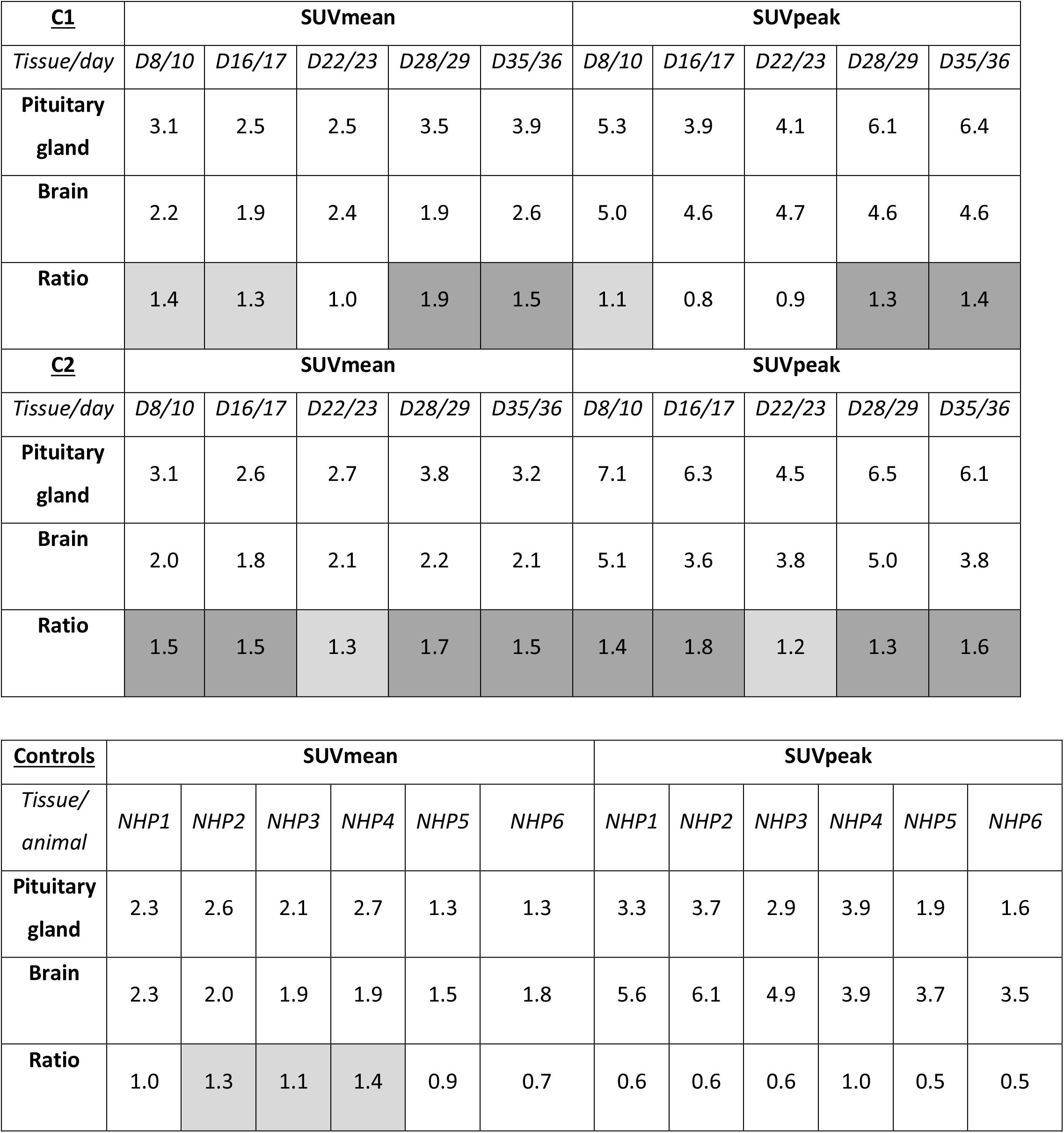
Quantitative PET-CT analysis of the macaque brains. The standardized uptake values (SUVs) of animals C1 and C2 are represented together with the SUVs of six non-infected control rhesus macaques. Of these animals the average and peak uptake are determined for the pituitary gland, and the brain minus the pituitary gland. By dividing these the pituitary gland/brain ratio is calculated. For the SUVmean values above 1.0 are defined as slightly increased (light grey) and increased (dark grey) when equal or above 1.5. For the SUVpeak values above 1.0 are demarcated as slightly increased and above 1.2 as increased.

